# Identification of Novel Fusion Genes in Pediatric B-ALL patients Using Whole Transcriptome Sequencing

**DOI:** 10.1101/2025.09.23.677992

**Authors:** Osheen Sajjad, Ali Amar, Abu Bakar, Muhammad Asif Naveed, Ayman Ali Mohammed Alameen, Saba Khaliq, Saqib Mahmood

## Abstract

**Background:** Fusion genes (FGs), serving as diagnostic, prognostic, and therapeutic markers, are key molecular aberrations in acute leukemia. They are also essential for risk stratification and measurable residual disease (MRD) monitoring. This study aimed to characterize the distribution of FGs using whole transcriptome sequencing (WTS).

**Materials and Methods:** A total of 12 newly diagnosed treatment naive pediatric B-ALL cases from a local tertiary care hospital were enrolled in this study. Following the nucleic acids isolation procedures, the RNA sequencing was done for 12 B-ALL patients to find the fusion genes.

**Results:** In the present cohort of 12 pediatric B-ALL patients, 19 high-confidence in-frame gene fusion events were identified involving 29 unique partner genes. The commonly reported sub-type defining rearrangements in B-ALL, including ETV6-RUNX1, TCF3-PBX1 and BCR-ABL1, were found in 8.3 % of the patients whereas the rearrangements in commonly prevalent genes in B-ALL like PAX5, ABL1 and ATXN3 were also found in 8.3 % of the patients with different unreported partner genes but reported earlier in various studies i.e. PAX-ETV6 (8.3%), ABL1-SNX2 (8.3%) and CMC2-ATXN3.

**Conclusion:** The present work expands the scope of fusions in pediatric B-ALL by revealing unreported and domain-retaining fusions, as well as co-occurring rearranged fusions with possible combinatorial outcomes. By introducing WTS into clinical workflows, the genetic classification can be done more accurately and new pathogenic drivers missed by traditional methods can be identified, highlighting the increased significance of transcriptomic profiling in the diagnosis, prognosis, and personalized therapy of leukemia.

## Introduction

Fusion genes (FGs) play a central role in acute leukaemia and as a founder event during leukemogenesis, as well as a disease-selective, diagnostic, prognostic, risk-stratification, and monitoring of MRD (1). The WHO classification has defined many fusion genes since 2001 as core diagnostic entities, screening and longitudinal monitoring of them are now routine parts of acute leukaemia management(2).

The most frequent subtype of ALL is B-cell ALL, which contains about 30 genetically defined subtypes that are caused by translocations, point mutations, and dysregulation of oncogenes and transcription factors, with consequences in cell cycle control, chromatin remodelling, survival signalling, and lineage commitment.(3, 4).

Among the hallmark cytogenetically visible rearrangements are t(12;21)(p13;q22)/ETV6-RUNX1, commonly associated with favorable prognosis; t(1;19)(q23;p13)/TCF3-PBX1, linked to CNS relapse; and the Philadelphia chromosome t(9;22)(q34;q11.2)/BCR-ABL1, found more frequently in adults and associated with poor outcome. KMT2A (MLL) rearrangements such as t(4;11)(q21;q23) are also prevalent in infant ALL and portend a dismal prognosis (5).

Subtypes characterized by cryptic fusions identified by transcriptome profiling include: ETV6-RUNX1-like and DUX4-rearranged ALL (∼8%), ZNF384 (∼5%) and MEF2D (∼7%) fusions denote other subgroups of B-ALL, with different prognoses and treatment responses.(5)

Kinase-driven leukaemia such as Ph-ALL and Ph-like ALL include ABL-class, JAK2 or PDGFRB fusions, particularly in adults, and show therapeutic potential with tyrosine kinase inhibitors.(6).

Functional categorization of FGs reveals that many act as transcription factors (e.g., RUNX1, ETV6), chromatin remodelers (e.g., KMT2A), or signaling molecules (e.g., ABL1, JAK2), highlighting their role in key leukemogenic pathways (7).

The fusion gene landscape defined by whole-transcriptome sequencing (WTS) is still incompletely resolved although multiple genetically defined subtypes of pediatric B-ALL are recognized. With respect to the structure, pathway context, and potential clinical relevance, in particular, cryptic and rare, domain-retaining fusions, as well as co-occurring rearrangements, remain under-characterized. Moreover, studies rarely apply standardized classification frameworks-combining caller confidence, pathogenicity tiering, and fusion-gene family (FG-FM) assignment-to enable consistent interpretation across cohorts. These gaps were addressed by using WTS to profile newly diagnosed pediatric B-ALL, systematically catalogued known and unreported fusions, evaluated domain retention and protein–protein interaction network context, and organized events using a tiering and FG-FM scheme to prioritize the ones with putative diagnostic, prognostic, or therapeutic significance.

### Subjects and Methods

#### Patients

From November 2021 to June 2022, 12 cases of paediatric B-ALL were recruited in Jinnah Hospital, Lahore, and diagnosis was done through flow cytometry, as per established protocols (8). Treatment-naïve patients aged ≤12 years of both genders were recruited at diagnosis, and excluded patients with congenital syndromes or genetic disorders. An informed written Consent by parents or their guardians were received prior to enrolment. The study protocol was approved by the Ethical Review Committee (No: UHS/EAPC-22/ERC/3) and the Advanced Studies and Research Board (No: UHS/Education/126-22/4352) of the University of Health Sciences, Lahore, and complied with the Declaration of Helsinki.

#### Sample Preparation

A minimum of 3-5 ml of peripheral blood containing ≥80% of blasts was harvested, enriched by Ficoll density centrifugation, viability assessed via Giemsa staining and stored at −20°C.

Isolation of genomic DNA was performed with the help of Quick-DNA™ kit and total RNA with Trizol. The quality of RNA was evaluated by Nanodrop and high-quality RNA was used to produce cDNA and sequence.

Complementary DNA (cDNA) synthesis was performed with the Maxima First Strand cDNA Synthesis Kit (Thermo Fisher Scientific, USA) using 500 ng to 1 µg of total RNA, according to the manufacturer’s protocol. The synthesized cDNA was stored at –20°C.

#### Library Preparation and Whole Transcriptome Sequencing (WTS)

Samples with adequate RNA quality were processed for RNA sequencing. Library preparation was performed using the NEBNext® Ultra™ Directional RNA Library Prep Kit for Illumina® (New England BioLabs) with ≥1 μg total RNA input. Sequencing was conducted using the NovaSeq 6000 platform (Illumina, USA) with 150 bp paired-end reads, yielding an average of ∼50 million reads per sample. Image analysis, base calling, and quality control were performed using Illumina RTA v1.18.64 and Bcl2fastq v1.8.4. FASTQ files were generated for bioinformatic analysis. The raw data of RNA-seq generated in this study have been deposited in the NCBI Sequence Read Archive (SRA) under accession number **PRJNA1332557**. The data settings were kept for private access to peer review and will be made publicly available upon publication. All patient data have been fully anonymized and no personal identifiable information was included.

#### Fusion Gene Detection from WTS Data

Fusion transcript detection was performed using the Arriba algorithm (version 2.4.0). Arriba assigns confidence levels (low, medium, or high) based on read support and background noise modeling. Only in-frame fusion transcripts with high-confidence calls were considered positive, excluding those annotated as read-through or artifacts. Reciprocal fusion events were counted as a single fusion event. Fusion genes were labeled “ Unreported “ if they were not previously reported in major fusion gene databases, including the Atlas of Genetics and Cytogenetics in Oncology and Haematology, the Tumor Fusion Gene Data Portal, and Chimer DB.

#### Clinical and Laboratory Data Collection

Diagnosis and subtyping were recorded using demographic, clinical, and laboratory data of 12 B-ALL patients using a structured pro forma, included age, gender, consanguinity, family history of cancer, duration of symptoms, and clinical signs, hematologic parameters, and flow cytometry-based CD markers.

The baseline characteristics, clinical features, and hematologic findings are summarized in Tables 1–3 and illustrated in Figures 4.1–4.3.

#### Fusion Gene Landscape and Pathogenicity Evaluation

Fusion analysis had 19 distinct genes, which were grouped into Tier A (4 known pathogenic drivers (e.g., *BCR-ABL1*), Tier B (14 likely pathogenic with rare/unreported partners), and Tier C (1 of uncertain significance).

## Results

### Spectrum and Incidence of Fusion Genes in Pediatric B-ALL

In 12 pediatric B-ALL patients, 19 in-frame fusions and 29 genes were observed:

Tier A (4/18, 22.2%, BCR:ABL1, ETV6:RUNX1, TCF3:PBX1). Fusion SNX2::ABL1, which involves the known ABL1 kinase and aligns with the molecular features of Ph-like acute lymphoblastic leukemia (ALL) was Unreported.

Tier B (5/18, 27.8%, functional kinases/transcription factors). Examples include MEX3C::ZBTB16, PRKCB::COTL1, and STIM1::CSNK2A1.

Tier C (9/18, 50%, unreported and with uncertain significance). These fusions include REEP5::UBB, PCBP1::ARPC3, and CMC2::ATXN3.

The majority of patients (7/12, 58.3%) harboured a single fusion, while some patients (4/12, 33.3%) had two distinct fusions. One patient (1/12, 8.3%) was found to have three concurrent fusion events. The primary mode of breakpoint was translocations, which occurred in coding/splice regions (72.2%).

An analysis of fusion event co-occurrence across 12 patients revealed distinct patterns of multi-tier fusion presentations.

Three patients (25.0%) had both Tier A and Tier B fusions. One patient (8.3%) presented with concurrent Tier B and Tier C fusions. Another patient (8.3%) harbored fusions from all three tiers: Tier A, Tier B, and Tier C.

It is noteworthy that no patient exhibited a direct co-occurrence of Tier A and Tier C fusions without the presence of a Tier B fusion. This suggests a potential hierarchical or sequential acquisition of fusion types, where Tier B fusions may act as a link between the other two tiers. (Fig. 1).

**Figure 1:**
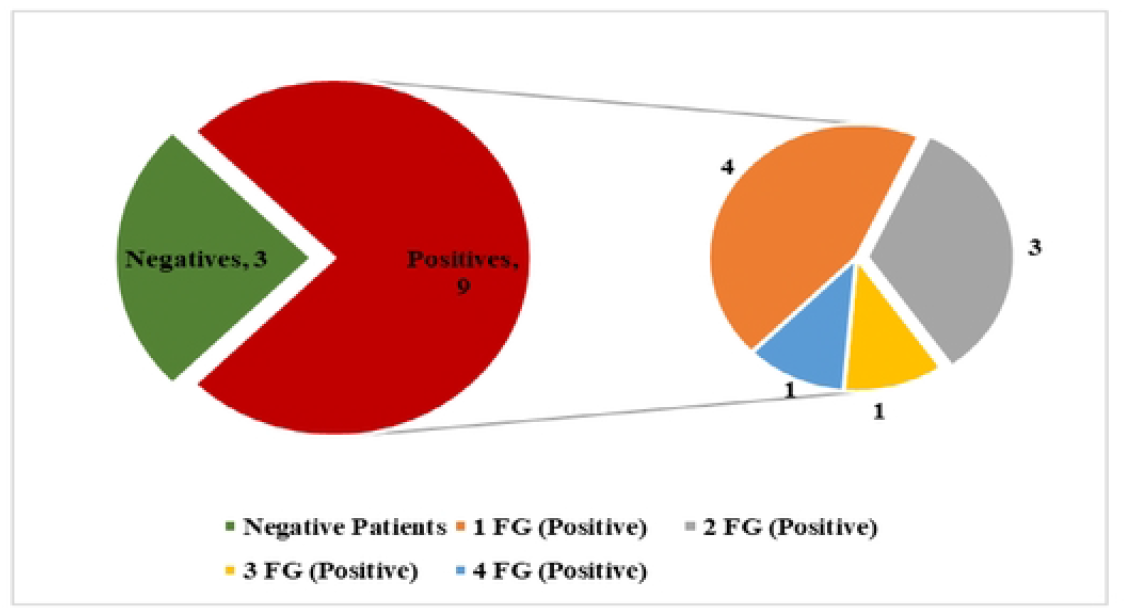
Distribution of Fusion Gene (FG) Positivity and Multiplicity in Pediatric B-ALL Patients. The left pie chart shows the proportion of patients with detectable fusion genes (FGs) (n=9, red) versus FG-negative patients (n=3, green). The right exploded pie chart highlights the number of distinct fusion genes identified per FG-positive patient: 4 patients harbored a single FG (orange), 3 had two FGs (gray), while 1 patient each showed three (yellow) and four (blue) FGs. This illustrates both the frequency and heterogeneity of FG events in the cohort.

### Fusion Gene Map of Pediatric B-ALL Cohort (Study Cohort)

To evaluate potential signaling convergence and oncogenic clusters, a curated set of 19 high-confidence fusion genes (FGs) identified in the study cohort was analyzed using a STRING-based protein–protein interaction network Figure 2.

**Figure 2:**
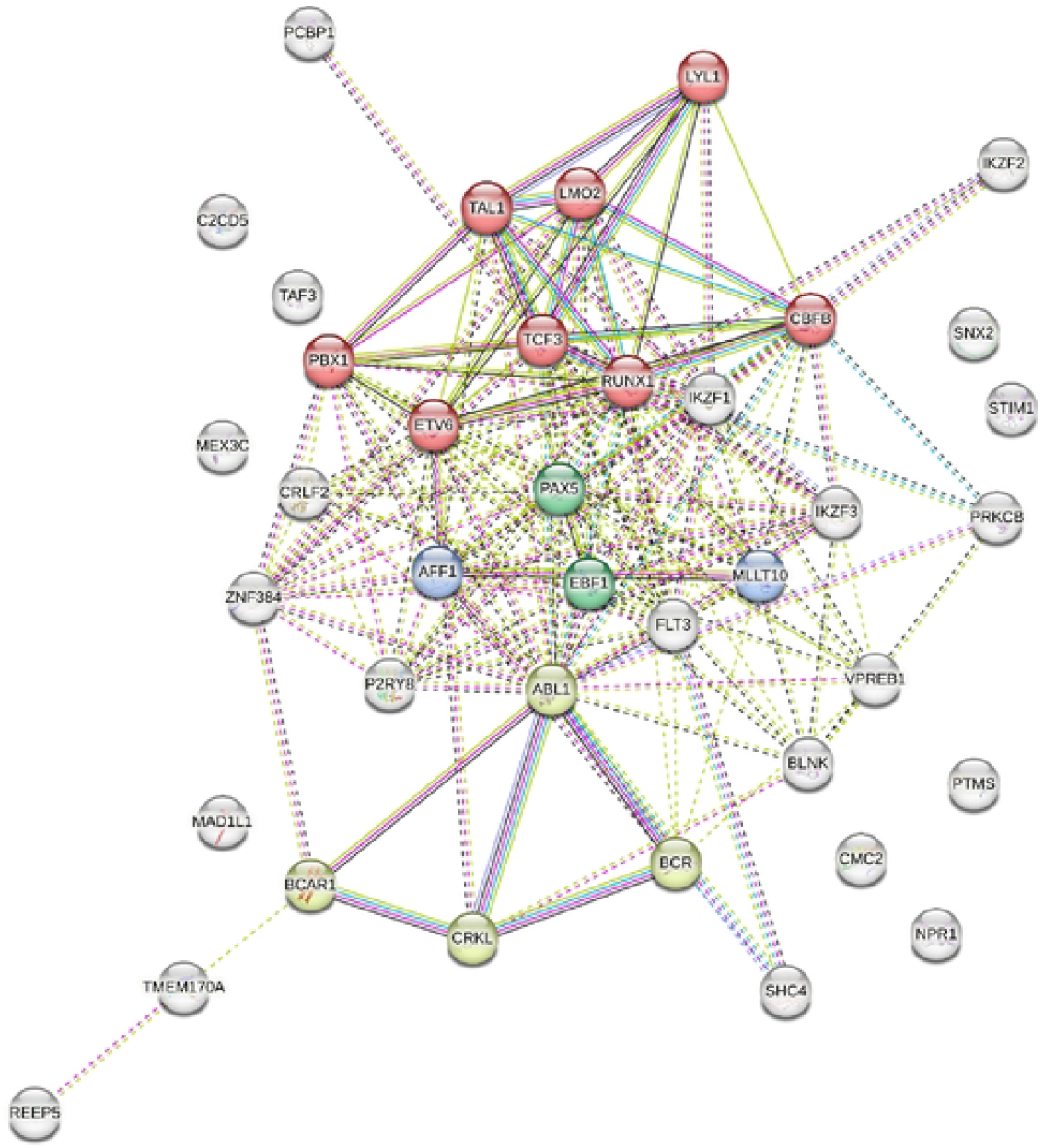
Fusion gene interaction network in pediatric B-ALL. STRING-based proteinprotein interaction network of fusion gene partners identified in pediatric B-ALL. Nodes represent gene products, and edges indicate predicted functional associations.

The interaction network showed that Tier A genes formed a highly connected cluster centered on well-known regulators of blood cell development (hematopoietic transcription factors) and signaling pathways (kinase regulators). Within this cluster, ABL1 and BCR acted as major hubs, linked to adaptor proteins such as CRKL and CBL, suggesting activation of downstream kinase signaling cascades. TCF3 also displayed strong connectivity, interacting with transcriptional partners including PBX1, PAX5, TAL1, and LMO2.

In contrast, Tier C genes (e.g., PTMS, NPR1, PCBP1) appeared at the periphery of the network, reflecting their limited known roles in hematopoiesis or leukemia biology.

Overall, this functional gene interaction map illustrates the diversity of fusion-driven pathways in pediatric B-ALL: Tier A fusions are deeply integrated into established oncogenic networks, Tier B fusions show partial or emerging links, while Tier C fusions remain poorly characterized.

### Protein kinase gene fusions

In our paediatric B-ALL cohort, kinase fusions were identified in 41.7% (5/12) of this cohort, with all gene pairs containing tyrosine kinases. The most common were ABL1 fusions (25%) and including ABL1-CRKL, ABL1-EBF1, and ABL1-ZNF384. FLT3-AFF1 and BCR-ABL1 fusions were each identified in one patient (8.3%). These effects of clinical significance reveal the importance of fusion screening to inform TKI-based treatment in paediatric B-ALL.

### Transcription factor gene fusions

Chromosomal translocations involving transcription factors are frequently seen in acute leukemia, and some of them have been used as genetic markers for leukemia classification because of their distinctive clinicopathological features and prognostic significance (9)

Current study showed in 50% (6/12) cases, 7 transcription factor fusions were detected, such as hallmark drivers such as TCF3::PBX1 and ETV6::RUNX1, important to B-ALL subtyping and risk stratification.

Most of the transcription factor fusions in our cohort involved core-binding factors (RUNX1, ETV6) and zinc-finger transcription factors (ZNF384), highlighting their recurrent role in B-ALL pathogenesis. The ZNF384 gene was found in fusion with both TCF3 and ABL1, aligning with its frequent involvement in pediatric B-ALL.

In our cohort, 2 epigenetic gene fusions involving MLLT10 and AFF1 were detected in 2 out of 12 cases (16.7%), both of which are implicated in histone modification and transcriptional regulation.

One case involved an AFF1-MLLT10 fusion, a rearrangement often associated with complex transcriptional dysregulation. The second case featured AFF1-FLT3, combining an epigenetic regulator with a tyrosine kinase, which may hold therapeutic implications across multiple regulatory pathways

### Unreported Domain-Retaining Fusion Genes as Potential Drivers in B-ALL

In this cohort, nineteen distinct, non-replicated fusion gene were defined widening the genomic spectrum of paediatric B-ALL, each occurring in a single patient, underscoring the individualized nature of these rearrangements as shown in the table 1. unreported combinations of many involved known hematopoietic regulators (e.g., ZNF384, IKZF1, PBX1, EBF1) indicate new mechanistic functions.

**Table 1:**
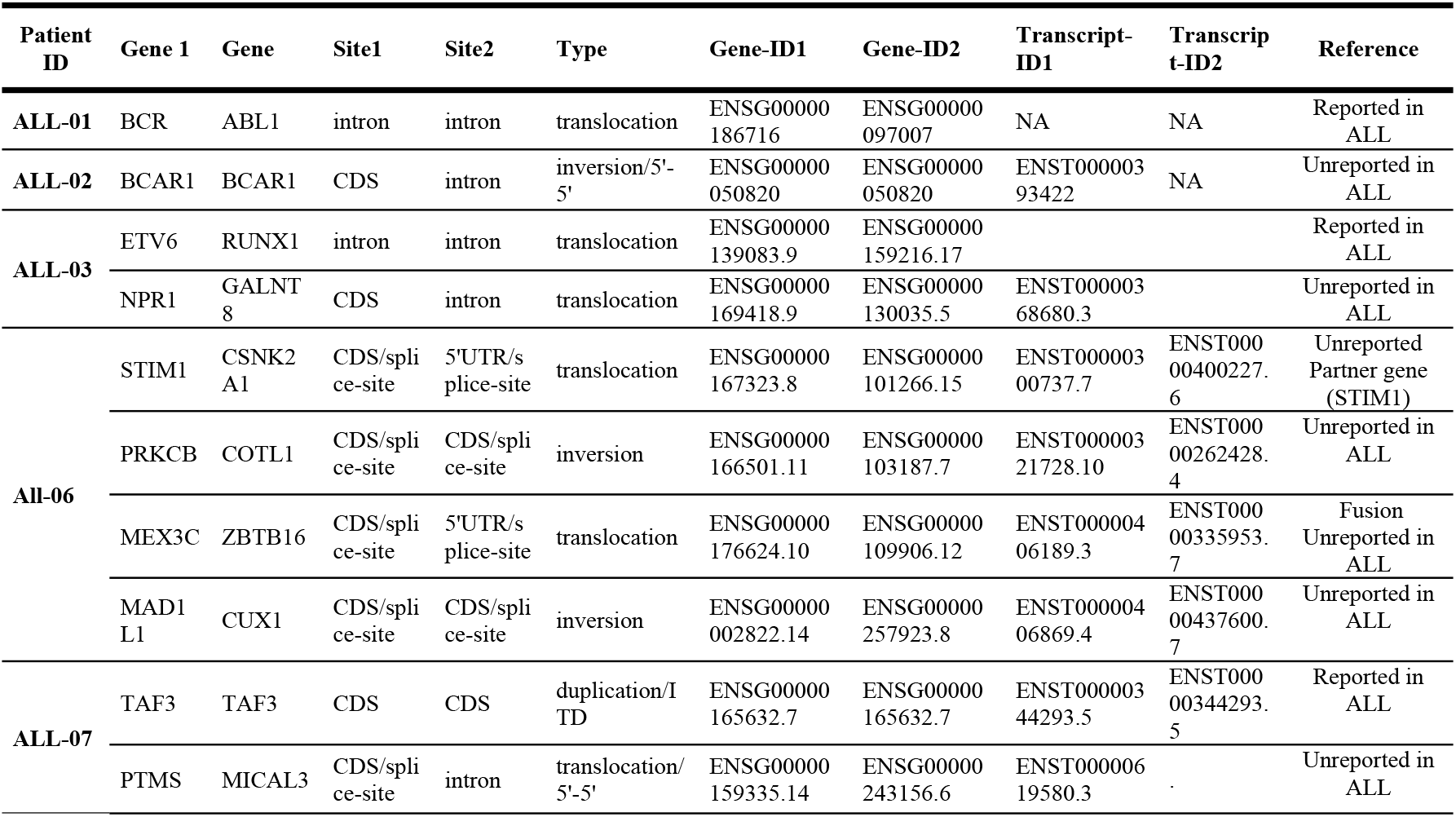
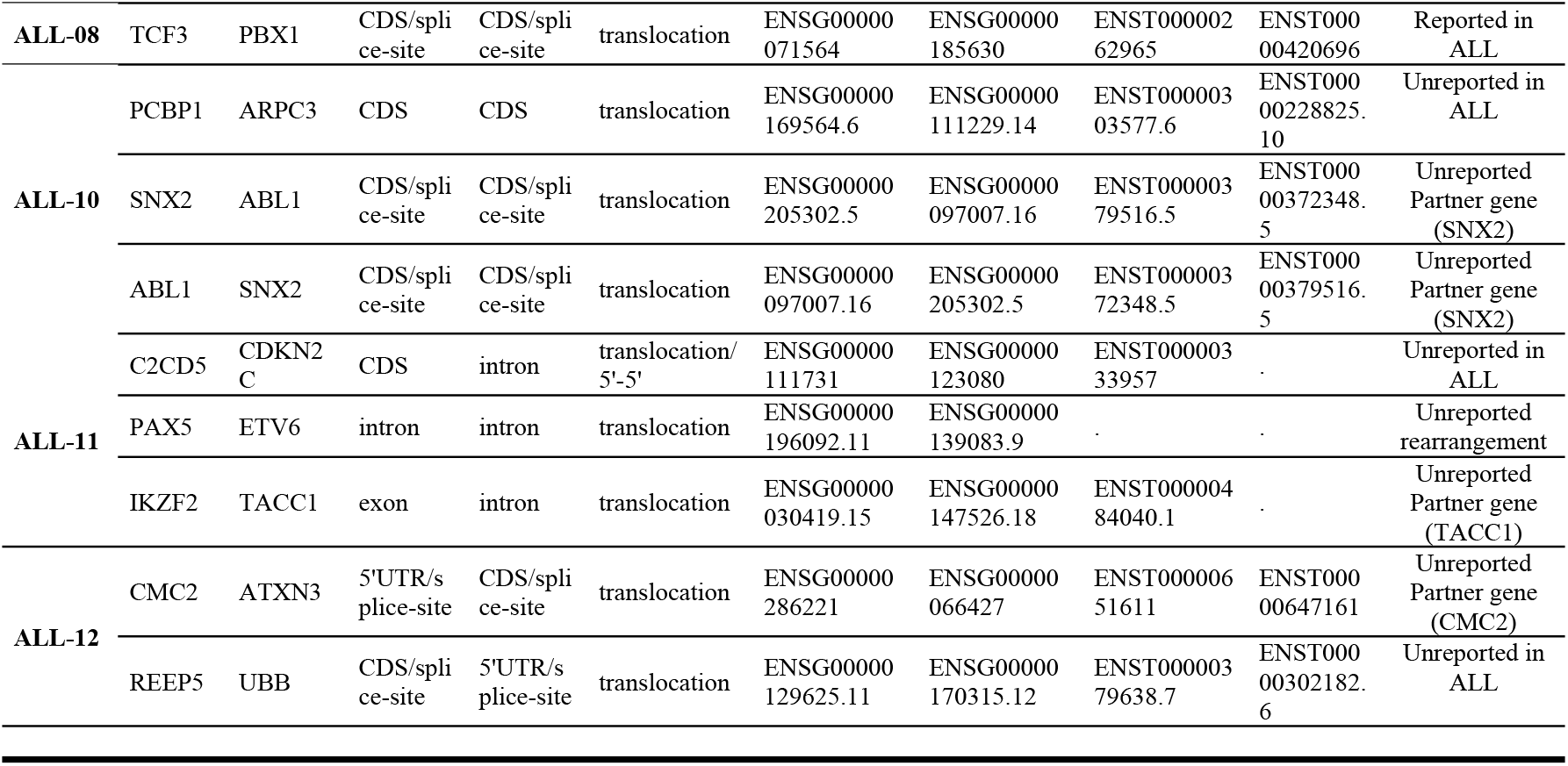
Fusion Transcripts found in the B Acute Lymphoblastic Leukemic Patients of the current study.

Significant new combinations comprised ZNF384:ABL1 and IKZF1:EBF1, which combine major regulators of signalling and B-cell differentiation. The majority of engaged coding-region breakpoints that had retained domains, which are indicative of functional effects.

Analysis at domain level identified five new fusions (BCAR1::BCAR1, NPR1::GALNT8, TAF3::TAF3, PCBP1::ARPC3, CMC2::ATXN3), all of which retained functional motifs related to signalling, cytoskeleton regulation, transcription or ubiquitin pathways, indicating the potential of leukemogenesis despite a lack of recurrence.

These fusions are illustrated below with exon-level breakpoint mapping and protein domain retention, highlighting their structural integrity and potential functional impact.

Inverted BCAR1-BCAR1 fusion on chromosome 16q23.1 with breakpoints at 16:75242553 and 16:75243902 is represented in figure 3. The top panel shows the gene structure and read coverage over each exon based on RNA-seq data, while the red dashed lines highlight the fusion junction. Though it is out-of-frame, it still retains intact SH3 domains, implying that it could cause changes in signalling and cytoskeletal organization, and circos plot (bottom left) confirms that it has an intra-chromosomal origin.

**Figure 3:**
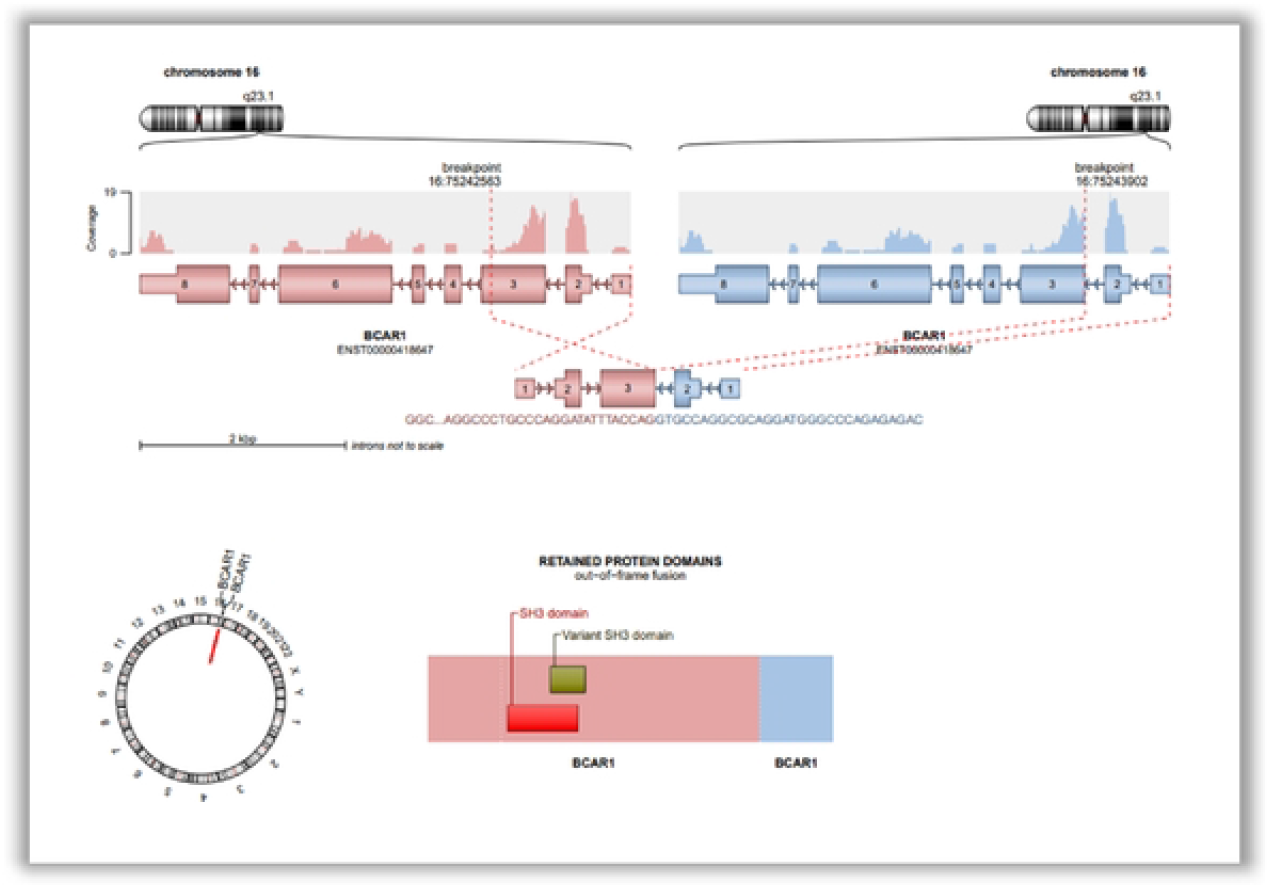
Fusion transcript arising from intra-chromosomai inversion of the BCAR1 gene.

In our cohort, BCAR1 (16q23.1) intragenic inversion created an out-of-frame *BCAR1-BCAR1* fusion that retained SH3 domains. (Fig. 3) Its contribution to adhesion and signalling has not been reported in leukemia, but its potential leukemogenic effect suggests it should be studied further.

Figure 4 gives the schematic illustration of the out-of-frame fusion between *NPR1* (chromosome 1q21.3) and *GALNT8* (chromosome 12p13.32). RNA sequencing coverage is displayed above the exon structures of both genes. Breakpoints are mapped to exon 1 of *NPR1* (ENST00000368660) and exon 15 of *GALNT8* (ENST00000648836). The circos plot (bottom left) shows inter-chromosomal rearrangement between chromosomes 1 and 12. The retained domain diagram (bottom right) highlights the preservation of the Receptor family ligand-binding region from NPR1. Introns are not drawn to scale.

**Figure 4:**
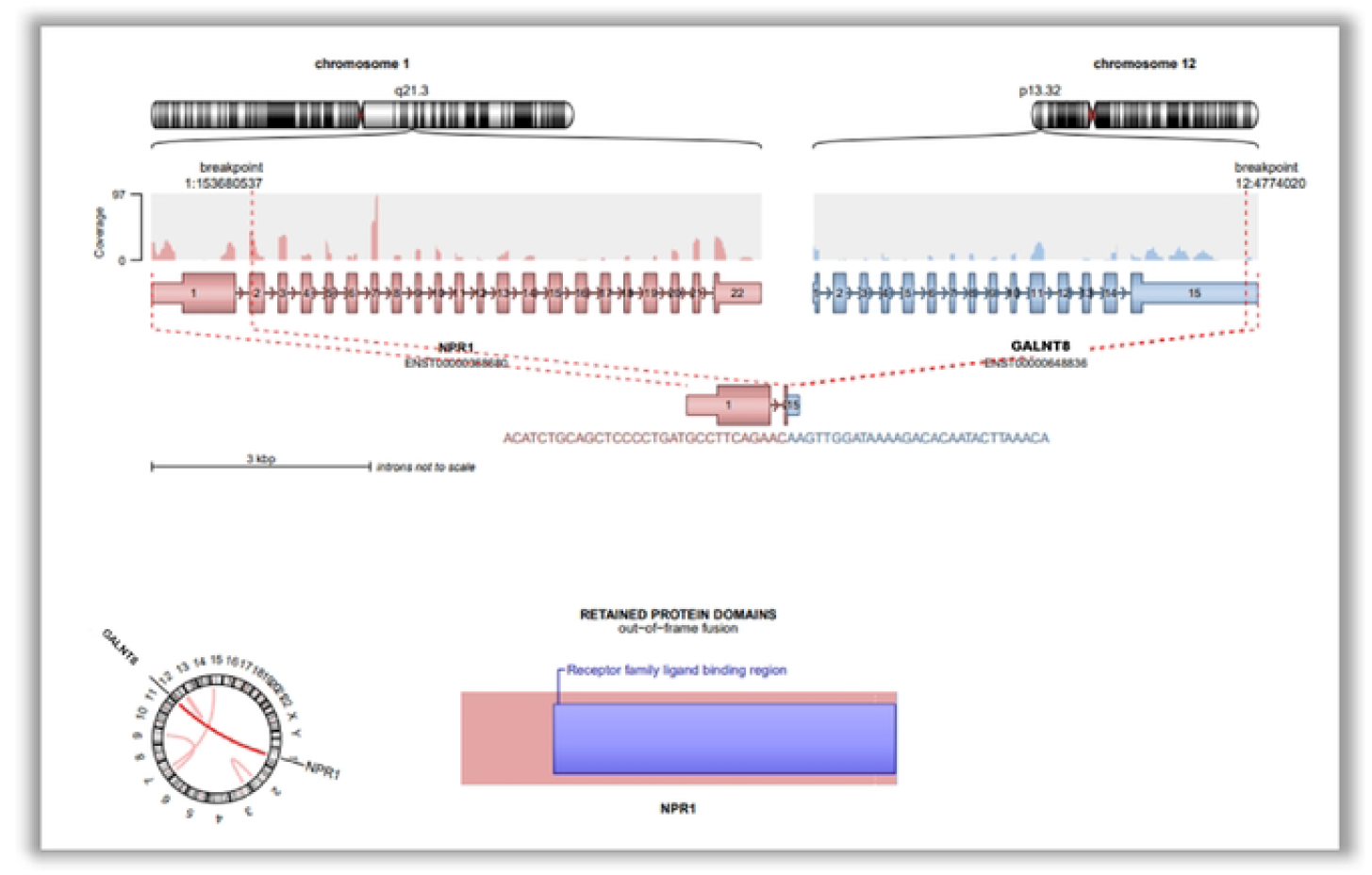
Structural representation and domain retention of the not reported NPR1::GALNT8 fusion gene.

The *NPR1::GALNT8* fusion is an inter-chromosomal rearrangement identified in a pediatric B-ALL case, which is Unreported in the previous literature. NPR1 encodes natriuretic peptide receptor A, which participates in cardiovascular signaling, apoptosis, and regulation of vascular tone, while *GALNT8* encodes a polypeptide N-acetylgalactosaminyltransferase involved in initiating mucin-type O-glycosylation. Though not previously linked in leukemia, both genes influence cell signaling and differentiation pathways. This out-of-frame fusion retains the receptor ligand-binding domain of *NPR1*, suggesting potential functional modulation in cellular communication. The combination of a signaling receptor and a glycosylation enzyme may have implications for leukemic cell surface dynamics and merits further investigation.

Figure 5 gives genomic illustration of a tandem duplication involving the TAF3 gene on chromosome 10p14, with breakpoints at positions 10:7967516 and 10:7965629. RNA-seq coverage tracks and exon-intron structures of the gene are shown with breakpoint-spanning reads. The fusion occurs between exon 3 and downstream exons of TAF3, generating a duplicated in-frame transcript. The circular plot (bottom left) depicts an intra-chromosomal rearrangement (green line = duplication). The domain map (bottom right) reveals retention of both the bromodomain-associated motif and the PHD finger, suggesting that chromatin-interacting functionality may be preserved.

**Figure 5:**
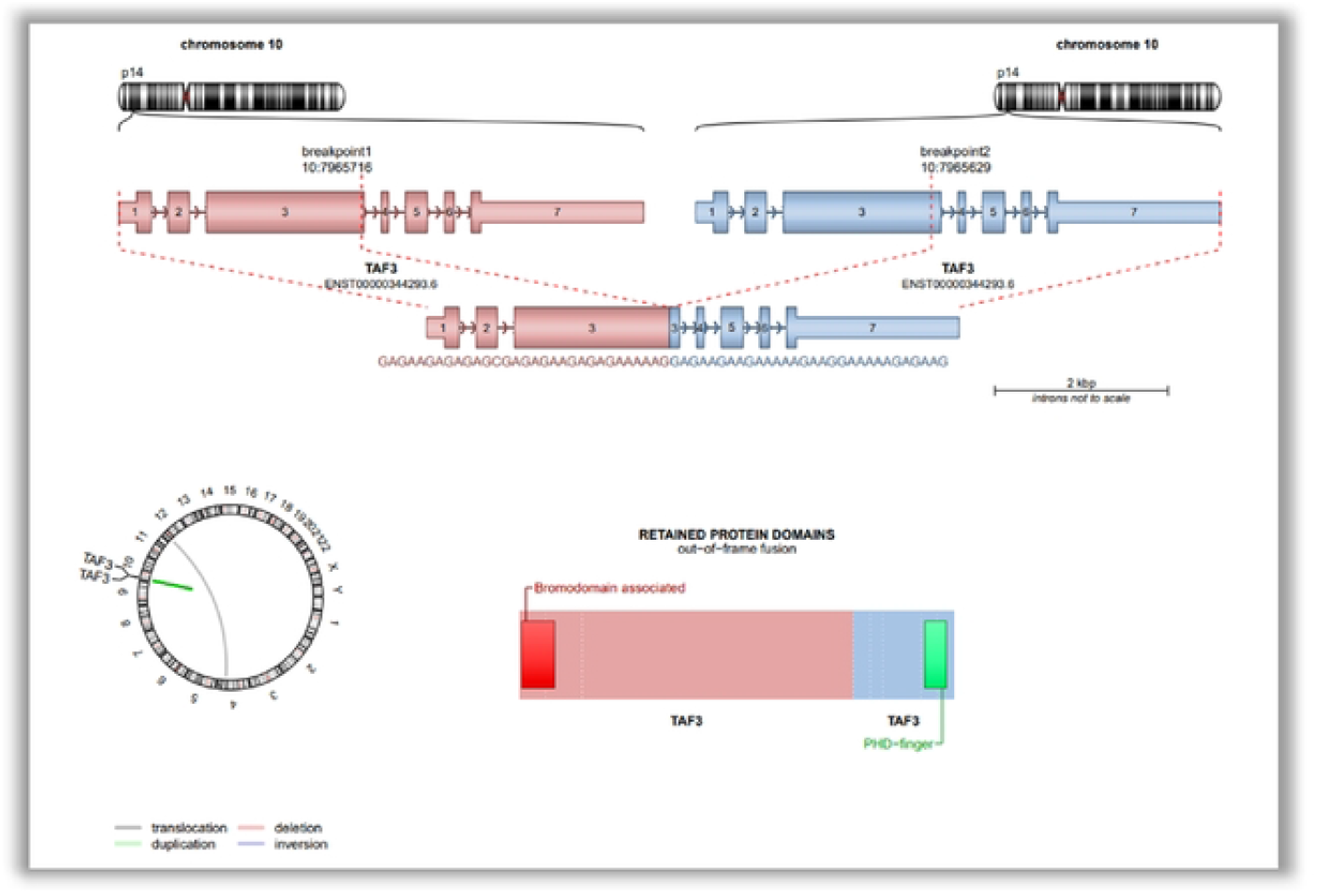
Structural depiction and domain retention of the TAF3::TAF3 tandem duplication fusion in B-ALL.

The TAF3::TAF3 fusion represents an unreported intra-chromosomal tandem duplication involving TAF3, a transcription initiation factor that binds histone H3K4me3 via its PHD finger and participates in chromatin remodeling and transcriptional regulation. This fusion preserves both major protein domains — the bromodomain-associated motif and the PHD-finger — despite being an out-of-frame event. These domains are crucial for recognizing chromatin modifications and may influence transcriptional programs relevant to leukemogenesis. While TAF3 has not been previously reported as a recurrent fusion gene in leukemia, its functional role in epigenetic control makes this duplicated transcript biologically intriguing and a candidate for further mechanistic exploration.

Figure 6 gives the schematic representation of an inter-chromosomal translocation involving PCBP1 (chromosome 2p13.3) and ARPC3 (chromosome 12q24.11). RNA-seq read coverage plots and exon structures are displayed with breakpoint positions and connecting fusion reads. The fusion joins exon 5 of PCBP1 to exon 5 of ARPC3, generating an out-of-frame transcript. The circular plot (bottom left) shows a translocation event linking the two chromosomes. The domain diagram (bottom right) indicates preservation of the KH (K Homology) RNA-binding domains from PCBP1 and the ARPC3 complex domain (21 kDa subunit) from ARPC3.

**Figure 6:**
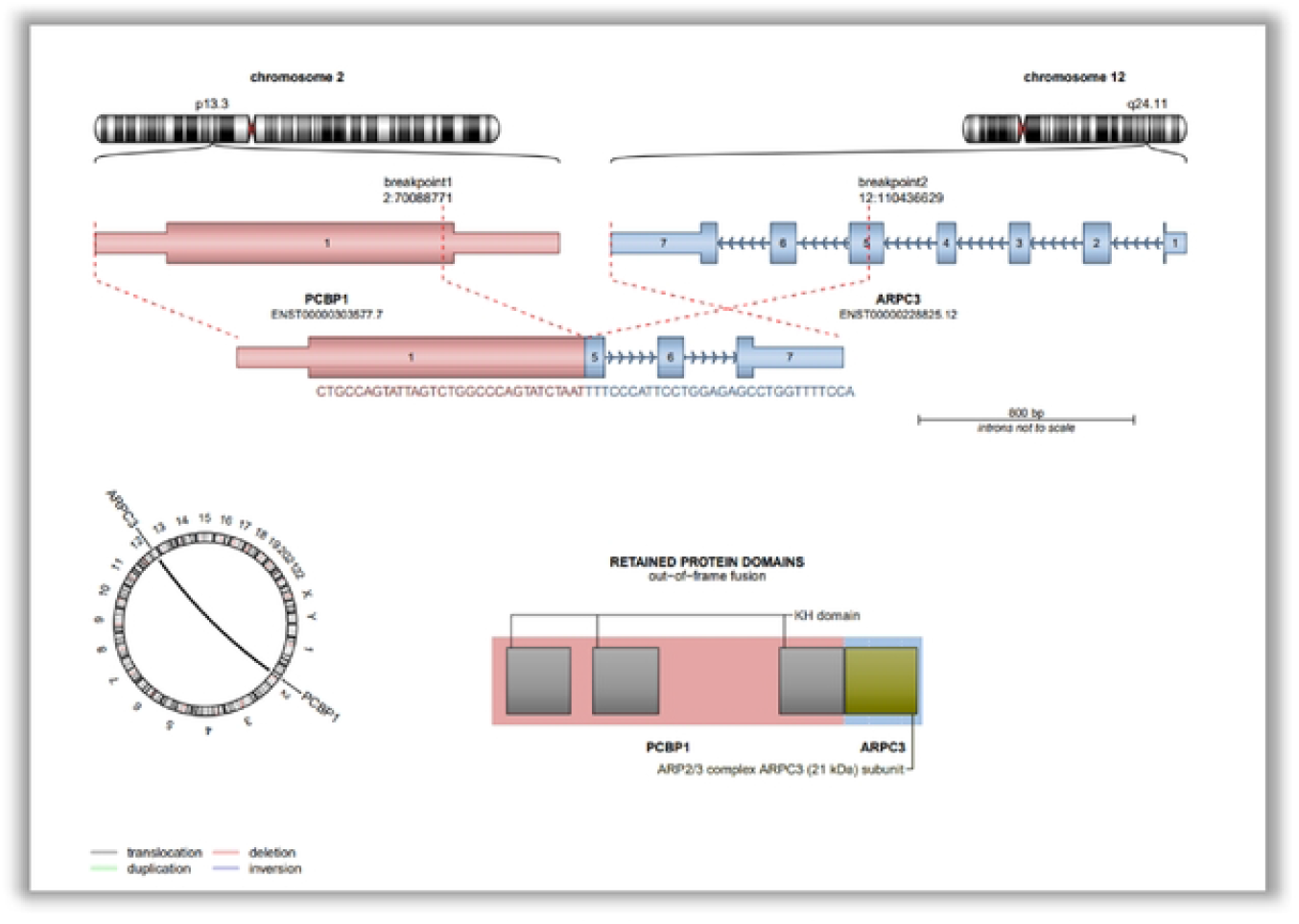
Inter-chromosomal fusion of PCBP1::ARPC3 showing domain retention in an out-of-frame transcript.

The PCBP1::ARPC3 fusion represents an unreported out-of-frame fusion event between PCBP1, a poly(C)-binding protein with multiple RNA-binding KH domains, and ARPC3, a core subunit of the actin-related protein 2/3 (ARP2/3) complex involved in cytoskeletal remodeling and cell motility. This inter-chromosomal rearrangement is biologically intriguing due to the combined involvement of post-transcriptional regulation (via PCBP1) and cytoskeletal signaling (via ARPC3). Although this specific fusion has not been previously reported in hematological malignancies, the preserved functional domains imply potential disruption of mRNA stabilization and actin dynamics. These features highlight the fusion’s relevance for future functional studies and its possible contribution to leukemogenic pathways. Figure 7 gives the schematic represents an unreported inter-chromosomal fusion between CMC2 (chromosome 16q23.2) and ATXN3 (chromosome 14q32.12). RNA-seq read coverage plots highlight breakpoints at exon 1 of CMC2 and exon 11 of ATXN3. Supporting reads were only observed on the ATXN3 side (n=2). A circular plot (bottom left) visualizes the translocation. Domain analysis (bottom right) shows multiple retained ubiquitin interaction motifs and a Josephin domain from ATXN3. No canonical protein domain was detected from CMC2 in the fusion transcript.

**Figure 7:**
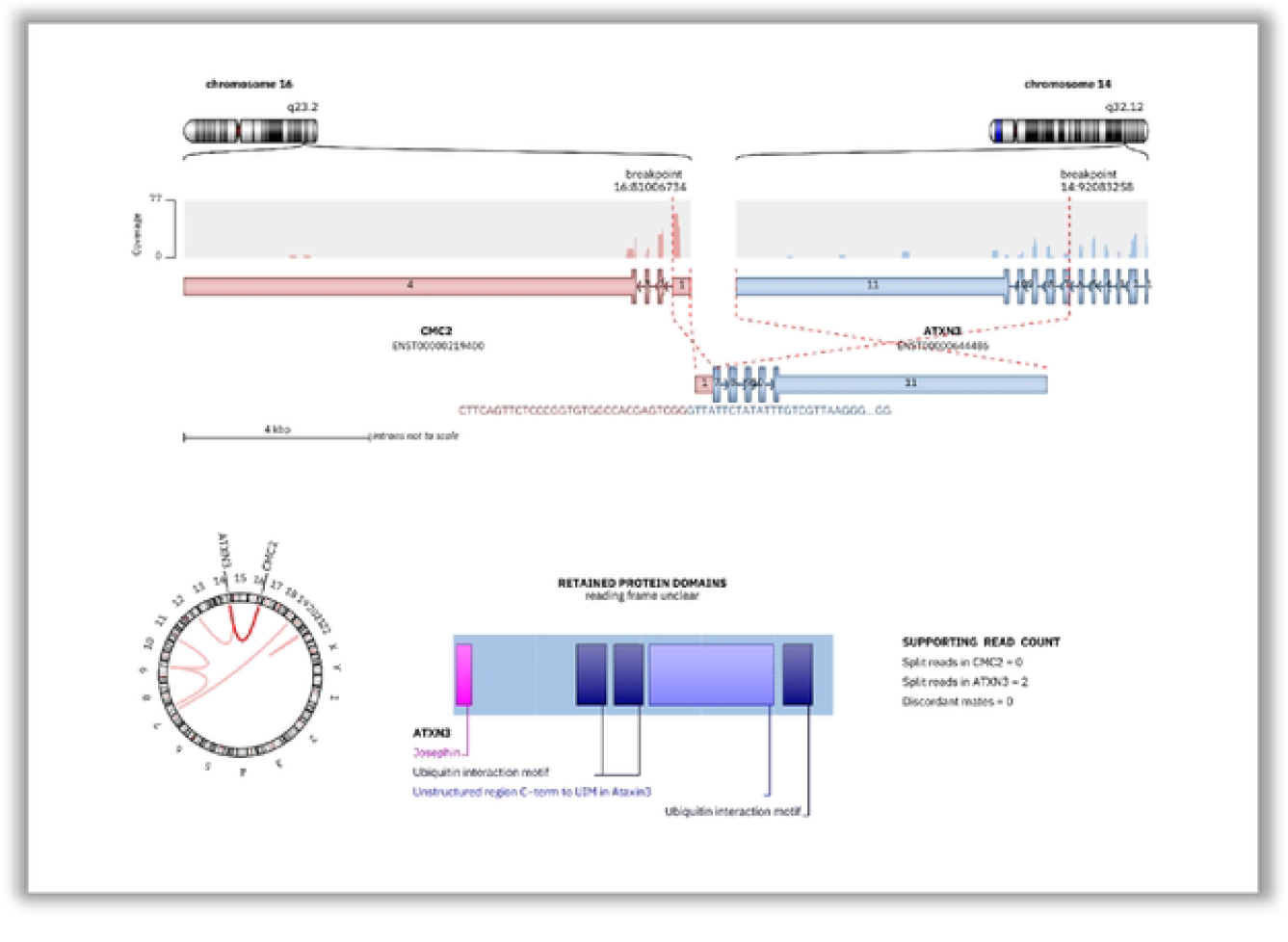
Inter -chromosomal CMC2::ATXN3 fusion involving retained ubiquitin-interaction domains in an unclear reading frame.

**Figure 8:**
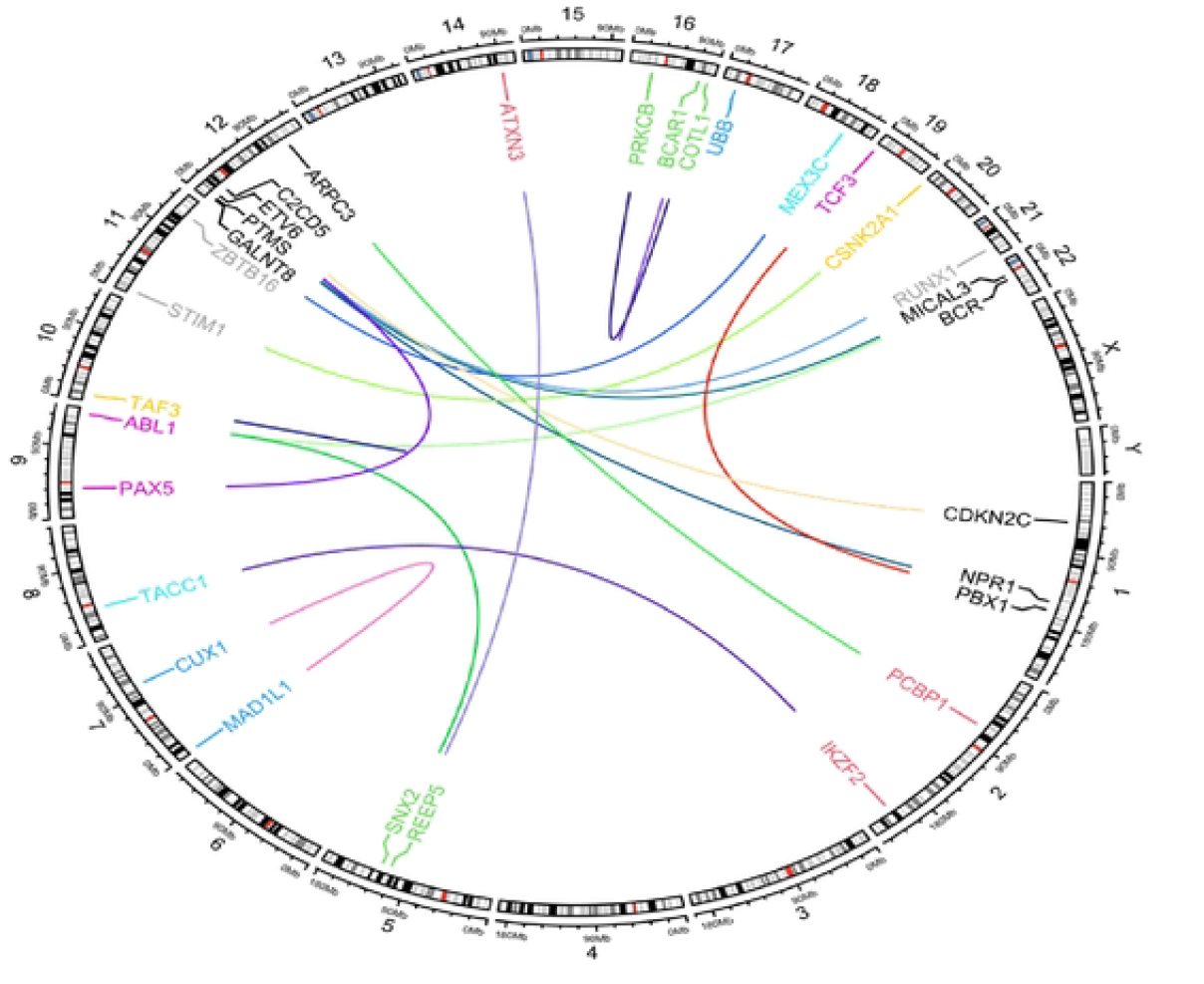
Circos plot showing chromosomal locations of fusion genes identified in pediatric B-ALL patients.

The CMC2::ATXN3 fusion represents a previously unreported rearrangement connecting two genes with distinct functions. CMC2 is involved in mitochondrial copper transport, while ATXN3 encodes a deubiquitinating enzyme implicated in neurodegenerative diseases. The preserved ubiquitin interaction motifs and Josephin domain in ATXN3 suggest partial retention of deubiquitinase functionality. However, the reading frame is unclear, and lack of reciprocal read support from CMC2 indicates potential instability or low expression of the fusion. Despite limited transcriptional evidence, the biological relevance of ubiquitin pathway involvement in leukemogenesis warrants further functional validation of this fusion, especially in the context of protein degradation and cell signalling.

### Classification of FGs according to FG-FMs

We classified the Tier A and Tier B fusion genes according to FG-FMs, defined as fusion gene models involving a recurrent “protagonist” gene with multiple distinct fusion partners. Among the 9 tier A and 7 tier B fusions (total 16), 6 fusions (37.5%) could be assigned to known FG-FMs, including ABL1-FM, ETV6-FM, ZBTB16-FM, CSNK2A1-FM, and CUX1-FM. These included hallmark rearrangements such as BCR::ABL1 and ETV6::RUNX1, as well as functionally related but less frequently reported fusions like SNX2::ABL1 and MEX3C::ZBTB16.

The remaining 10 tier A/B fusions (62.5%), such as PRKCB::COTL1, PAX5::ETV6, and STIM1::CSNK2A1, could not be definitively assigned to existing FG-FM families, although they involved known cancer-related genes. Most unclassified fusions occurred only once and represent rare or potentially unreported events.

These findings underscore the heterogeneity of fusion gene landscapes in pediatric B-ALL and suggest that even within a small cohort, a subset of fusions aligns with known FG-FM families, reinforcing their biological and clinical relevance.

### Chromosomal distribution and genomic architecture of fusion genes

To visualize the chromosomal locations and inter-chromosomal rearrangements of fusion genes (FGs) detected in our cohort, we constructed a genome-wide Circos plot (Fig.8). Each arc represents a gene fusion event connecting two genomic loci across chromosomes, with distinct colors denoting different fusion pairs. The outer ring represents human chromosomes labeled by their cytogenetic bands. Several prominent hotspots were identified on chromosomes 9 (hosting ABL1, PAX5, TAF3), 12 (ETV6, LMO2, CBFB), and 22 (BCR, RUNX1), which are well-known in leukemogenic translocations. Notably, intra-chromosomal and inter-chromosomal rearrangements were both observed, reflecting the heterogeneous mechanisms of gene fusion formation in B-ALL. The diversity in genomic loci and cross-chromosomal fusion connections underlines the structural complexity of leukemic genomes and supports the need for comprehensive sequencing approaches to capture clinically relevant fusions.

## Discussion

In this study, RNA sequencing (RNAseq) revealed fusion genes in approximately 61.8% of paediatric cases of B-ALL, in agreement with data worldwide, which revealed known and unreported events and highlights the value of transcriptome profiling.

Rearrangements defined as subtypes (ETV6-RUNX1, TCF3-PBX1, BCR-ABL1) were detected in 8.3% of the patients, which is noteworthy compared to Kimura et al. (2021), but rather low compared to the standard 20-25% prevalence of ETV6-RUNX1 in paediatric B-ALL (10, 11). B-ALL rearrangements, PAX5, ABL1 and ATXN3 were observed in 8.3% of Unreported partners, in line with previous findings including PAX5-ETV6 (8.3%), ABL1-SNX2 (8.3%), and CMC2-ATXN. PAX5–ETV6 fusions have been reported previously in ALL cohorts (12). The biological and diagnostic relevance of transcription factor fusions in child leukemia is underscored, which justifies their incorporation into diagnostic and prognostic systems.

Numerous new fusions, primarily between transcription factors, kinases, and chromatin remodelers, were identified in this cohort. Even though no KMT2A or EP300 fusions were identified in our cohort, AFF1 and MLLT10 fusions were observed, which indicates the use of chromatin remodelling genes in paediatric B-ALL and makes epigenetic-targeted therapies worth considering (13).

Recent findings indicate that chromatin-modifying and epigenetic genes are crucial cancer drivers, thus they are potential therapeutic targets. Approved HDAC and DNA methylation inhibitors confirm epigenetic therapies. Several new combinations such as ZNF384 -ABL1 and IKZF1 -EBF1 have been identified connecting important lymphoid regulators, and ZNF384 - ABL1 is a distinct pathogenic interaction (14, 15).

Domain-level analysis identified unreported fusions that preserve functional domains (involved in signal transduction, transcriptional regulation, and post-translational modification), including NPR1::GALNT8, TAF3::TAF3, PCBP1::ARPC3 and CMC2::ATXN3, indicating them as individualized disease drivers and possible biomarkers or treatment targets (16, 17, 18). These new events are patient selective, uncommon and could serve as drivers of B-ALL, highlighting the importance of transcriptome profiling in the discovery of diagnostic or therapeutic fusions.

Translocations were the most common source of fusion events detected in exonic, intronic, and untranslated regions by RNAseq. Particularly, it is notable that 35.7% were intra-chromosomal and many of them remained invisible to conventional cytogenetics, which underscores chromosomal instability in the evolution of leukemia. (19).

Interestingly, multiple fusion events were detected in several patients, including one case harbouring four distinct fusions; indicate sub clonal diversity or genomic instability. The interplay of co-occurring Tier A, B and C fusions can be cooperative, although the biology of such interaction remains to be investigated.

In line with other large-scale studies, our data identify RNAseq or WTS as the best method in the identification of fusion gene, where cryptic, rare, and variant events are revealed beyond the scope of multiplex RT-PCR. Its domain architecture solving capability as demonstrated by the new KMT2A fusions reinforces its utility in accurate diagnostics (20).

Not every fusion identified by sequencing is pathogenic, many of them are passenger events. Tier-based and FG-FM classification can be used to rank clinically relevant rearrangements. For example, PAX5::JAK2, although involving a major B-cell transcription factor, is more appropriately categorized under JAK2-driven leukemias due to its functional mimicry of BCR-ABL1-like disease (21).

From a clinical standpoint, several FGs identified in this cohorts such as BCR:ABL1, SNX2:ABL1 and new JAK2:CSNK2A1 fusions have therapeutic implications, and ETV6:RUNX1 and TCF3:PBX1 inform prognosis and risk stratification. These fusions are also used as MRD markers in treating patients (22, 23).

In conclusion, the fusion gene landscape in this study is expanded in pediatric B-ALL, uncovering unreported and concurrent events. The implementation of WTS into clinical practice allows a more accurate genetic classification and the development of this method in the direction of diagnosis, prognosis, and individual treatment. Although the mechanistic insights may be broadly relevant but these estimates should not be taken as globally representative without corroboration in independent, multi-regional cohorts

